# Phylogenetic Biogeography Inference Using Dynamic Paleogeography Models and Explicit Geographic Ranges

**DOI:** 10.1101/2023.11.16.567427

**Authors:** J. Salvador Arias

**Author notes:** Emai correspondence:. Laboratorio de Genética Evolutiva, Instituto de Biología Subtropical (CONICET-UnaM), Facultad de Ciencias Exactas, Quimicas y Naturales, Universidad Nacional de Misiones, Felix de Azara 1552, CP 3300, Posadas, Misiones, Argentina.

## Abstract

Most popular methods of phylogenetic biogeography discard the spatial component of geographic distributions, dividing Earth into a handful of predefined areas. Other methods use explicit geographic ranges, but unfortunately, these methods assume a static Earth, ignoring the effects of plate tectonics and the changes in the landscape. To address this limitation, I propose a method that uses explicit geographic ranges and incorporates a plate motion model and a paleolandscape model directly derived from the models used by geologists in their tectonic and paleogeographic reconstructions. The underlying geographic model is a high-resolution pixelation of a spherical Earth. Biogeographic inference is based on diffusion, approximates the effects of the landscape, uses a time-stratified model to take into account the geographic changes, and directly integrates over all probable histories. By using a simplified stochastic mapping algorithm, it is possible to infer the ancestral locations as well as the distance and speed traveled by the ancestral lineages. For illustration, I applied the method to an empirical phylogeny of the Sapindaceae plants. This example shows that methods based on explicit geographic data, coupled with high-resolution paleogeographic models, can provide detailed reconstructions of the ancestral areas but also include inferences about the probable dispersal paths and traveling speed across the taxon history that are not possible with current methods based on predefined areas.

## INTRODUCTION

The main objective of phylogenetic biogeography is to infer the evolution of geographic ranges within a clade based on its phylogenetic relationships and the observed geographic locations of its terminals (Brundin 1965, 1966, 1972; Nelson 1969; Ronquist and Sanmartín 2011). Most current methods model geographic range data using predefined areas. These approaches include character mapping (Bremer 1992; Ronquist 1994; Clark et al. 2008; Landis 2016; Gunnell et al. 2018; Landis et al. 2021), dispersal-vicariance analysis (DIVA, Ronquist 1997), and the dispersal-extinction-cladogenesis model (DEC, Ree et al. 2005; Ree and Smith 2008; Matzke 2014) and its derivatives (Goldberg et al. 2011; Webb and Ree 2012; Landis et al. 2022). However, the predefined areas approach is problematic (Ree and Sanmartín 2009; Arias et al. 2011; Landis et al. 2013; Quintero et al. 2015; Arias 2017; O’Donovan et al. 2018). Even detailed models have a small number of areas (up to 25 by Landis 2016); the usual analysis uses between four and twelve areas. As a consequence, most predefined areas have large surfaces, clumping together the ranges of many species, even if their ranges are allopatric. The boundaries of these predefined areas are usually poorly defined, and in most cases, no primary data is provided for area assignment, hindering the repeatability of the research. Predefined areas are usually defined for each different paper, making it difficult to compare the results of different studies. While there are some *ad hoc* solutions proposed for these problems (e.g., constructing a distance-based dispersal matrix, Webb and Ree 2012; Landis 2016; Landis et al. 2022; or an automated process to assign predefined areas, Töpel et al. 2016), these do not address the fundamental problem: the geographic data of the terminals is discarded.

Using explicit range data, such as georeferenced specimen locations or geographic range maps, offers an alternative to the predefined area model. There are two main approaches that use explicit geographic range data. The first approach is based on the ideas of DIVA and DEC, in which an evolutionary model of dispersal and extinction is used to infer changes along branches, and widespread ranges are assigned to internal nodes. These methods are implemented using parsimony (Arias et al. 2011; Arias 2017; see also Hovenkamp 1997, 2001) or a probabilistic evolutionary model (Landis et al. 2013). The second approach is more similar to character mapping, in which a diffusion process models the evolution along the branches without taking extinction into account. These models were initially developed for intra-specific phylogeography (Lemmon and Lemmon 2008; Lemey et al. 2010; Bouckaert et al. 2012; Pybus et al. 2012; Bouckaert 2016; Louca 2021) and then extended to model inter-specific conventional phylogenetic biogeography (Nylinder et al. 2014; Quintero et al. 2015; O’Donovan et al. 2018). These models assign a single point to the ancestor instead of a widespread range distribution, which speeds up the computing time and directly calculates the likelihood of any geographic location (only observed locations are used in Arias et al. 2011; Arias 2017; additional locations must be explicitly included by the user in Landis et al. 2013).

There are several developments to make the diffusion model more realistic, including using a spherical Earth (Bouckaert 2016; Louca 2021) or taking into account the effects of landscape (Bouckaert et al. 2012).

Although explicit geography methods provide an important advance in modeling the evolution of geographic ranges, they fall short in other aspects. In particular, one of the main advantages of the predefined area methods is the incorporation of a paleogeographic model (Ree et al. 2005; Ree and Smith 2008; Webb and Ree 2012; Landis 2016). All methods using explicit geographic ranges lack this critical element of inference, so they effectively assume that Earth’s geography is static or that the effects of the dynamic nature of the Earth in the evolution of ranges are minimal (O’Donovan et al. 2018). There is another limitation of methods based on explicit geographic ranges: they rely on data augmentation in internal nodes (Lemmon and Lemmon 2008; Landis et al. 2013; Quintero et al. 2015; Bouckaert 2016; O’Donovan et al. 2018), then Monte Carlo sampling is required to approximate the integral over all possible histories, which makes most analyses slow.

The objective of this paper is to introduce a phylogenetic biogeography method that uses explicit geographic ranges, uses a spherical Earth, is based on diffusion, and incorporates a dynamic paleogeography model as part of its inference machinery. The method accounts for changes in the location of continents (plate tectonics) and includes changes in the landscape (e.g., marine transgressions). Different from predefined area methods, these paleogeographic models are the same models used by geologists for tectonic reconstructions (e.g., Müller et al. 2019, 2022). Taking advantage of the geographic data model, this method provides direct integration across all possible histories using the pruning algorithm (Felsenstein 1981). Although developed for inter-specific phylogenetic biogeography in deep time, this method is also applicable for infra-species phylogeography analyses. Thanks to its reliance on models of high spatial and temporal resolution, this method will allow researchers to gain a more detailed understanding of the spatial component (Quintero et al. 2015; O’Donovan et al. 2018) and the geographical context of the range evolution than is not possible using current predefined area methods.

## MODEL COMPONENTS

An overview of the input data, the elements used for the reconstruction, and the products of the method is given in Figure 1.

**Figure 1.**
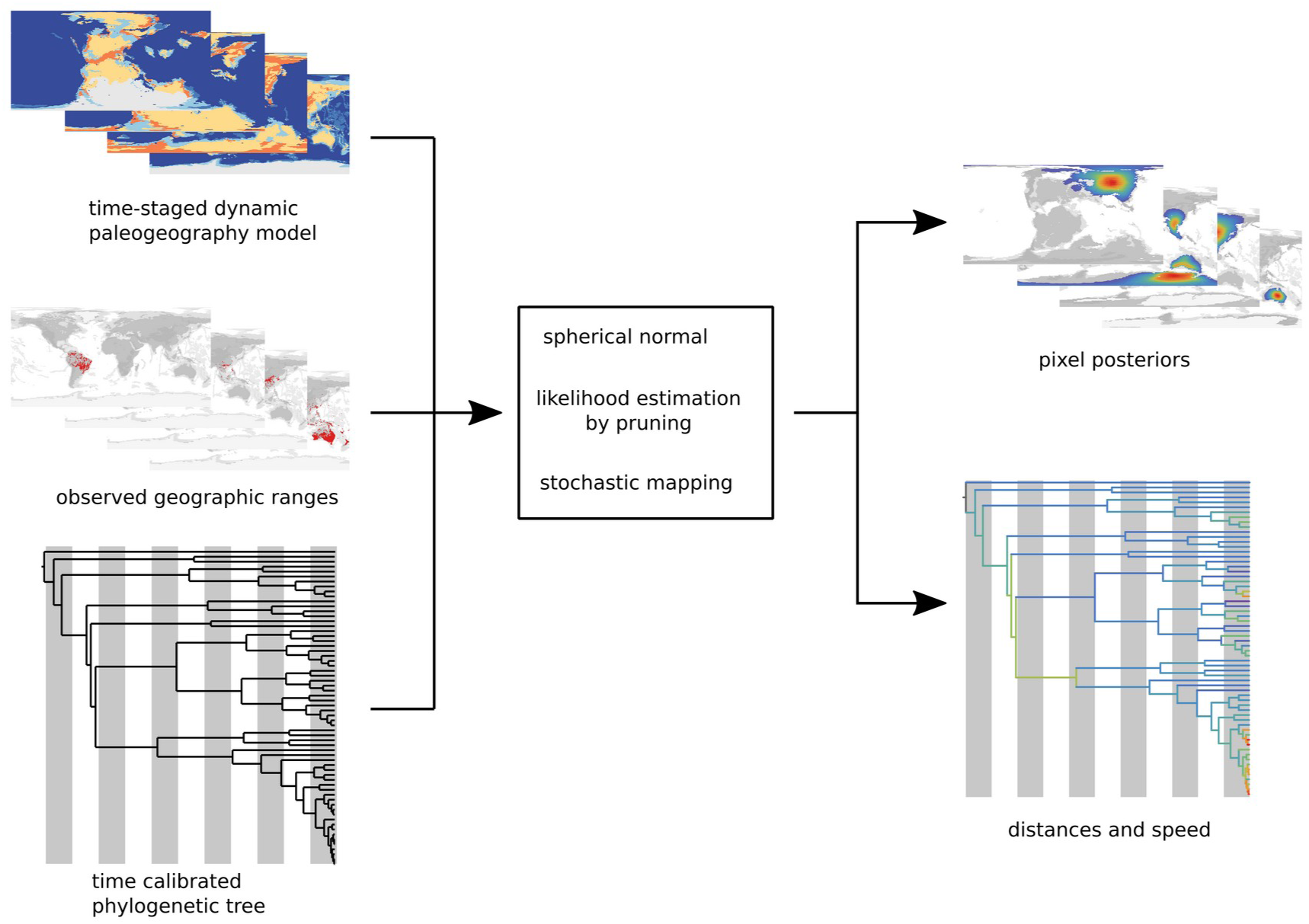
A sketch overview of the method presented in this paper. The basic input data for a phylogenetic biogeography analysis used here are the dynamic paleogeography model, the distribution data, and a time-calibrated phylogenetic tree (left). This data is used in the reconstruction phase (center) by modeling diffusion with a spherical normal and optimizing it by the Felsenstein pruning algorithm, and then stochastic mapping is used to produce the main output of the method: the pixel posteriors and the speed and distance of the reconstructions (right).

### Geographic Data Model

Geographic data is typically modeled in two ways: the vector model and the raster model (Fotheringham et al. 2000; Neteler and Mitasova 2008). In the vector model, spatial objects are represented by points, lines, and polygons. While this data model is continuous, it is typically used to represent discrete units such as the contours of continents, islands, or rivers. It has been applied in phylogenetic biogeography for diffusion-based methods (Lemmon and Lemmon 2008; Lemey et al. 2010; Bouckaert et al. 2012; Pybus et al. 2012; Nylinder et al. 2014; Quintero et al. 2015; Bouckaert 2016; O’Donovan et al. 2018; Louca 2021). The raster model represents geographic space as a matrix of pixels. While this data model is discrete, it is commonly used to represent continuous variables, such as climate data or digital elevation models. The raster model has been applied in phylogenetic biogeography (Arias et al. 2011; Arias 2017) as well as in other fields of biogeography, such as distribution modeling (Phillips et al. 2006) or identifying areas of endemism (Szumik and Goloboff 2004). Although either the vector or raster model can represent any geographic data, the decision to use one or the other is generally based on ease of implementation. Here, I use a raster model because it is straightforward to use for continuous fields, which are critical for the implementation of the pruning algorithm.

Most vector and raster models in biogeography use an equirectangular projection (e.g., Lemey et al. 2010; Arias et al. 2011; Pybus et al. 2012; Quintero et al. 2015; Arias 2017), which assumes a flat, cylindrical Earth and produces high distortions outside the equatorial region. Some vector model implementations use a spherical Earth (Bouckaert et al. 2012; O’Donovan et al. 2018; Louca 2021). Here, I use a raster model based on equal area partitioning (Saff and Kuijlaars 1997) that takes into account the sphericity of the Earth. This pixelation divides Earth into latitude rings of equal size, and each ring is divided according to its circumference, using the size of a pixel in the equatorial ring. Each pole has its own pixel, and the starting point is offset by a half pixel in odd rings. This pixelation is easy to implement, and pixels can be located quickly. It also ensures that the area of pixels is more or less the same (in fact, the total number of pixels can be calculated as *pix = eq^2^ ÷ π*, where *eq* is the number of pixels at the equator; this formula is the same as the area of a sphere using pixel units) and has a good isotropy, meaning that on average, each pixel has the same number of neighbors at a given distance (supp. Fig. 1).

To indicate the resolution of a pixelation, I use the notation e<NUMBER>, where <NUMBER> represents the number of pixels in the equatorial ring. For example, e360 denotes a pixelation with 360 pixels at the Equator.

The spatial resolution of the pixelation is dependent on the computational power available. As the number of pixels increases exponentially with resolution, any increase in resolution significantly impacts memory usage and computation speed. For example, an e120 pixelation has 4,586 pixels, while an e360 pixelation has 41,258 pixels. To achieve higher resolutions with a manageable number of pixels, it may be useful to prohibit certain pixels. For example, prohibiting pixels in deep ocean areas on an e360 pixelation can reduce the number of pixels to approximately 18,500.

### Dynamic Paleogeography Models

To model continental drift, geologists build plate motion models. These models are based on Euler’s theorem and involve a sequence of rotations along different axes of rotation on a sphere, which can be used to infer the location of a tectonic feature at a particular time (these models are also known as rotation models; for an introduction, see Cox and Hart 1986). These models are usually vectorial and continuous in time (e.g., Müller et al. 2019, 2022; Merdith et al. 2021). Here, to reduce computational time, plate motion models are pixelated and divided into time stages in which the geography is assumed to be static, as is done in DEC and other methods (Ree et al. 2005; Ree and Smith 2008; Webb and Ree 2012; Bielejec et al. 2014; Landis 2016). Under the model used here, each pixel at the current time is associated with a tectonic element and an assigned age. Then the plate motion model is used to determine the location of a pixel in an older stage while preserving its identity. However, due to the discrete nature of the pixelation, there can be instances where the location equivalent to a pixel after rotation can be in two or more pixels. As tectonic features are moved independently of any other feature, two or more different pixels can be moved to the same location after rotation. As tectonic features have a particular age, no pixel is assigned if the time stage is older than the pixel age. These issues are addressed with specific procedures during the reconstruction (see below in the RECONSTRUCTION section).

Although plate tectonics is a major driver of ancient geography, landscapes are also dynamic; for example, a location that is currently on land may have been part of an epicontinental sea in the past. This can be represented using a paleolandscape or paleogeography model. Typically, these models are represented as raster files or vector polygons and divided into discrete time stages (e.g., Cao et al. 2017; Kocsis and Scotese 2021). In the method presented here, the paleolandscape model is pixelated using the same spatial and temporal resolution as the plate motion model, and each pixel at every time stage is assigned a landscape identifier (e.g., shallow sea, lowlands, ice sheets). It is important to note that while the paleolandscape model stores pixel locations, it does not track pixel identities over time (that is done by the plate motion model). Keeping both models separated is a flexible solution, as the same plate model can be coupled with different paleolandscape models.

The temporal resolution of dynamic paleogeography models is a matter of practicality: it must be fine enough to make the static Earth assumption valid yet long enough to minimize computations over long branches. Moreover, the models should be available for the expected time stages. Fortunately, recent developments in free and open software such as GPlates (Müller et al. 2018) and publicly available datasets, like those found at https://www.earthbyte.org/category/resources/data-models/global-regional-plate-motion-models/, make it easier than ever to create and edit these models. While biologists can build these models by themselves (e.g., Bolotov et al. 2022), I think that it can be better seen as an opportunity for interdisciplinary collaboration, as paleogeographic models must be based on large amounts of geological evidence (e.g., Müller et al. 2022). Probably the most common situation is using an already defined dynamic paleogeography model. With this in mind, I created a GitHub repository (https://github.com/js-arias/geomodels) with ready-to-use dynamic paleogeography models as well as instructions for importing the user’s own GPlates model.

### Phylogeny and Terminal Geographic Ranges

The method presented here requires a fully dichotomous, time-calibrated phylogenetic tree. Although the method can be adapted to polytomic trees, I will not attempt to do it here. In cases where a lineage intersects with one or more time stages as defined in the dynamic paleogeography model, it is divided into distinct time slices, and inter-nodes are inserted between these divisions as is done in DEC and other methods (Ree et al. 2005; Ree and Smith 2008; Webb and Ree 2012; Bielejec et al. 2014; Landis 2016).

Each terminal must include an explicit geographic range. This geographic range can take the form of georeferenced specimen locations or continuous range maps. In the case of specimen locations, the localities are transformed into a presence-absence pixelation. For continuous range maps, each pixel within the range map is assigned a value; in a simple range map, all pixels have the same value, whereas distribution models allow for different values per pixel. During setup, regardless of the source, the values stored in terminal ranges are scaled so that the sum of all pixel values is equal to 1.0 (Bouckaert et al. 2012).

### Diffusion Model

The geographic range of a species can be seen as a collection of particles that move randomly across the Earth’s surface. Under that approach, the movement of these particles can be modeled using a diffusion process.

In phylogenetic biogeography, most diffusion models are based on the assumption of a flat Earth (Lemmon and Lemmon 2008; Lemey et al. 2010; Bouckaert et al. 2012; Pybus et al. 2012; Nylinder et al. 2014; Quintero et al. 2015). Under this assumption, diffusion is only the traditional normal distribution scaled over time, allowing speed-ups such as Gibbs sampling (Lemey et al. 2010; Quintero et al. 2015) or just storing the means and variances (Pybus et al. 2012; Nylinder et al. 2014). Under a flat Earth model, it is also possible to use different diffusion parameters for latitude and longitude (Lemey et al. 2010; Pybus et al. 2012; Quintero et al. 2015). But distances calculated using latitude and longitude as flat coordinates are neither Euclidean nor spherical, with distortion particularly pronounced near the poles. O’Donovan et al. (2018) used a 3D model of the Earth and computed a normal distribution for each axis with the same diffusion parameter using the Euclidean distance between point coordinates. While O’Donovan et al. (2018) conditioned start and end points at the Earth’s surface, they used a flat measure that ignores Earth sphericity, and its distortion relative to great circle distances increases with geographic distances. Moreover, as the movement is integrated over a 3D space, impossible paths in the Earth’s interior will have higher likelihoods than paths over the Earth’s surface.

An explicit spherical normal is a more preferable function for diffusion over the spherical surface of the Earth (Brillinger 1997; Bouckaert 2016; Louca 2021). Several approximations to the spherical normal have been proposed (e.g., Pennec 2006; Ghosh et al. 2012; Hauberg 2018), and for this paper I use the approximation given by Hauberg (2018):

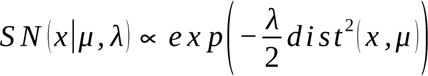

where *x* is a point on the sphere, μ is the mean point, λ is the concentration parameter in units of radians^-2^ (analogous to the κ parameter of the von Mises-Fisher distribution or the precision in the flat normal distribution), *exp* is the exponential function, and *dist* is the great circle distance between two points. One of the drawbacks of the spherical normal is its isotropy. Although anisotropic formulations exist (Hauberg 2018), their axes must follow a geodesic line (a great circle) on the sphere, meaning that the concentration matrix is uniquely associated with a particular mean point; therefore, it is not possible to use the same matrix when the mean point is rotated to a new position (Hauberg 2018).

Hauberg (2018) provides a function to scale the spherical normal at any point, but for the purpose of this paper, and taking advantage of equal area partitioning of the pixelation, a numerical integration over all pixels is used to produce a discrete approximation of the spherical normal.

## RECONSTRUCTION

The main issue with event-based methods is that they treat widespread ranges as the units of inference (Ronquist 1994; Ree et al. 2005; Landis et al. 2013), which results in an exponential increase in the number of potential ancestral ranges with the number of pixels (or predefined areas). To obtain the reconstructions of the ancestors, all these widespread ranges are integrated out, so their identity is discarded, and the result is the posterior probability of each individual pixel (Landis et al. 2013). An alternative approach is to focus directly on a single pixel (or location) as the unit of inference. The goal is to calculate the probability that a pixel in a node gives rise to any of the observed pixels in each descendant of the node (Lemmon and Lemmon 2008; Lemey et al. 2010; Bouckaert et al. 2012; Pybus et al. 2012; Nylinder et al. 2014; Quintero et al. 2015; Bouckaert 2016; O’Donovan et al. 2018) instead of the probability of the whole ancestral range.

### Calculating the Likelihood

Ideally, distances and the landscape can be taken into account through a numerical integration of the spherical normal over small time segments (Bouckaert et al. 2012). However, this approach can be computationally expensive and is limited to relatively small problems in time and space (Bouckaert et al. 2012). Therefore, most methods ignore landscape effects in the diffusion process and allow particles to move freely across the entire geography (Lemmon and Lemmon 2008; Bouckaert et al. 2012; Pybus et al. 2012; Quintero et al. 2015; Bouckaert 2016; O’Donovan et al. 2018). Here, the landscape is only taken into account to condition the probability of arrival at a particular point (Bouckaert et al. 2012).

To model the diffusion from an ancestral point to a descendant point on a branch segment, a probability function *f* is defined as:

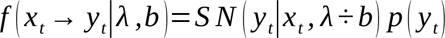

where *SN* is either the continuous or discrete spherical normal; *x* and *y* are points at time stage *t*; *x* is the point at the beginning of the branch segment, and *y* is the point at the end of the segment; *b* is the length in time of the branch segment, which is used to scale the λ parameter with time (Ghosh et al. 2012; Bouckaert 2016), just as is done in the planar diffusion; and *p* is the prior of the arrival point at time stage *t*. The priors of the points are defined using the pixel values in the landscape model.

Then the likelihood of a phylogenetic tree *T* with *n* nodes, including terminals, split nodes, and inter-nodes in each time segment of a branch, given a set of terminal locations *D* and a λ parameter, using *f* with the continuous spherical normal, is (Bouckaert 2016):

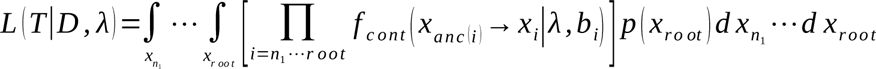

that is the sum of all possible histories from any point in the root to the observed points in the terminals (Lemey et al. 2010; Bouckaert et al. 2012; Pybus et al. 2012; O’Donovan et al. 2018). This integral is analytically intractable (Lemmon and Lemmon 2008; Bouckaert 2016). Therefore, a numerical approximation is necessary. The fastest alternative is to use independent contrast (Louca 2021) but geographic locations on the ancestral nodes, one of the main objectives of the analysis, are lost. The common approach to keeping the inference of ancestral pixels is to sample a single point from each terminal (Bouckaert et al. 2012; Nylinder et al. 2014; O’Donovan et al. 2018) and augment the data by assigning a single point to each internal node, either to maximize the likelihood (Lemmon and Lemmon 2008; Bouckaert 2016) or to perform an MCMC integration (Lemey et al. 2010; Bouckaert et al. 2012; Quintero et al. 2015; Bouckaert 2016; O’Donovan et al. 2018). Quintero et al. (2015) were able to skip the terminal sampling and use the whole geographic range of terminals, but continued to use the MCMC step for the augmented data in the internal nodes.

But if the problem is seen through the lens of a raster model, full numerical integration can be done by transforming the integrals of an infinite number of points into sums of pixels, using *f* with the discrete spherical normal:

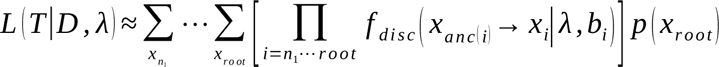

this formulation has an identical structure to the problem solved by Felsenstein (1981) with the pruning algorithm as recursive multiplications of conditional likelihoods.

The conditional likelihood of a pixel in the terminals is just the scaled value used for the terminal ranges, either a presence-absence pixelation or a range map, so that the sum of all pixel values is equal to 1.0 (Bouckaert et al. 2012; Quintero et al. 2015).

For an internode *m*, the conditional likelihood for the pixel *x* at the time stage *t* is (Ree and Smith 2008):

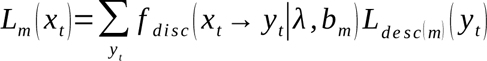

At inter-nodes, there is a change in the time stage, so the pixel locations are updated by applying the rotation model to move the conditional likelihood of a pixel from its previous location (at a younger age) to a potentially different location in the new time stage (at an older age). If multiple pixels are moved to a single destination, the maximum value is retained (i.e., likelihoods are not summed). If the source pixel has multiple destinations, the same value is assigned to all pixels (i.e., likelihoods are not divided). This procedure is made to maintain the smoothness of the ancestral likelihood field. If there is no destination pixel, the likelihood vanishes.

In the splits, the conditional likelihoods of both descendants of *n* are multiplied:

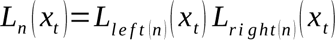

At the root, the full likelihood is the sum of the conditional likelihoods of all pixels at the root multiplied by the prior of the pixels at the root stage:

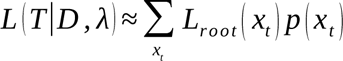

### Estimation of **λ** Parameter

With the likelihood function, it is possible to estimate the value of λ using maximum likelihood or Bayesian estimation. The likelihood function seems to be quite smooth, so it can be maximized by a simple hill-climbing algorithm. There is no known conjugate prior for the spherical normal (Hauberg 2018), so Gibbs sampling cannot be implemented, but as only a single parameter is estimated instead of using an MCMC algorithm, the Bayesian posterior can be computed directly by a numerical integration of the product of the likelihood and the prior.

### Estimation of Ancestral Pixels

I use a stochastic version of the demarginalization procedure of Yang et al. (1995) to estimate the pixel posterior probabilities at nodes. This approach allows for additional estimates, for example, of the average distance traveled by a particle (O’Donovan et al. 2018). This procedure is equivalent to a reduced version of stochastic mapping (Nielsen 2002; Dupin et al. 2017) and is labeled as such in the remainder of this paper. The posterior probability of a pixel at the root of the tree is the final likelihood of the pixel divided by the likelihood of the whole reconstruction:

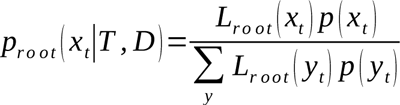

For each internode *m*, the probability of a destination *y* is the probability of reaching that pixel from the ancestral pixel *x* multiplied by the conditional likelihood of *y* at the end of the segment *m*, scaled by all possible arrivals:

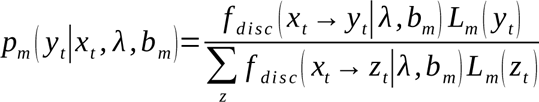

When a time stage changes, the location of a source pixel is changed using the plate motion model. As mentioned earlier, it is possible that the source pixel is represented by two or more destination pixels in the younger time stage. To solve this problem, a single destination pixel is randomly selected using the priors of the destination pixels in the younger time stage. At splits, both descendants inherit the same particle location.

This procedure is repeated multiple times, resulting in a collection of pixel locations and particle trips. The frequency with which a pixel is selected provides an approximation of the posterior probability for that pixel at a node during a given time stage. As trips are stored, the traveled distances of each trip can be calculated. As particles are moved with the plate motion model as the time stage changes, but the distance measure is taken within a single time stage, this approach only measures the distance traveled by the particle for reasons different from plate motion.

## IMPLEMENTATION

The method described in this paper is already implemented. The source code is written in the *Go* programming language and is available at https://github.com/js-arias/phygeo. This implementation takes advantage of the Go concurrency model and the current multiprocessor architecture to run the pruning algorithm and the stochastic mapping in parallel. There are two additional shortcuts worth mentioning. First, as the spherical normal is isotropic and the neighborhood of each pixel is also nearly isotropic, it is possible to precalculate the discrete normal density of any pixel in a latitude ring (i.e., a discrete spherical normal centered in the north pole) and then create a lookup table for the probability at each distance. Second, it will be possible to build a precalculated distance matrix for all pixels. This matrix requires more memory resources (for example, the distance matrix for an e360 pixelation takes about 1.4 GB), but because it removes the calculation of the great circle distances in the reconstruction phase, it produces a noticeable speedup in the calculations.

When the branch leading to a terminal taxon is small and the value of λ is large, it is possible that the full likelihood of far-away pixels cannot be calculated in a fast way. In such cases, only the maximum value will be stored (i.e., the most probable path from the source pixel to any destination); this will provide a reasonable approximation of the full likelihood (Bouckaert 2016) for mostly irrelevant pixels (i.e., pixels far away from the destination in a low diffusivity scenario).

The implementation has a reasonable speed for the size of the current datasets. In the empirical data set (see below), calculating the likelihood of a given λ parameter (including 1000 stochastic mappings) takes about 60 minutes on a 10-plus-year-old i5 machine with four cores at 2.9 GHz. The same data set runs in about 9 minutes on a modern M1 MacBook with eight cores at 3.2 GHz. Then estimating the maximum value of λ takes less than a day, while the approximate Bayesian estimation (see below) takes less than a week.

## BIOGEOGRAPHY OF SAPINDACEAE

To demonstrate the use of the model presented here in an empirical case, I made a biogeographic analysis of the Sapindaceae, a world-wide distributed, mostly tropical, family of angiosperms in the order Sapindales. The phylogeny of this group (Buerki et al. 2009, 2011, 2013, 2021; Joyce et al. 2023) provides a fair representation of many current analyses, in which the whole group is well sampled, but it is incomplete at the species level. While there are previous biogeographic analyses for the group (Buerki et al. 2011, 2013), they are based on predefined area methods, and the phylogenetic relationships are slightly different from the tree used here (Joyce et al. 2023).

## Materials and Methods

### Paleogeographic model

The paleogeographic model uses an e360 pixelation. The plate motion model is the Müller et al. (2022) model, with time stages separated by 5 Myr (for a total of 22 time stages from present to 105 Ma). The paleolandscape model is the Cao et al. (2017) model, rotated with the Müller et al. (2022) model. The full paleogeographic model (which extends to 540 Ma) is available at https://github.com/js-arias/gm-muller-2022. In the previous analyses (Buerki et al. 2011, 2013), they used seven areas in four time stages, so the current paleogeographic model represents an increase of 2600 times in spatial resolution and five times in temporal resolution. The prior probability for pixels was set at 1.00 for emerged land (either low or highlands), 0.005 for shallow sea on continental shelves, and 0.001 for oceanic plateaus and continental ice sheets.

### Phylogenetic tree

The phylogenetic tree was extracted from the Sapindaceae branch of Joyce et al. (2023) phylogenomic analysis of Sapindales. As the original publication does not provide a machine-readable file, the relationships and ages were extracted manually from their figure 4a. The phylogeny was augmented with a few terminals from Buerki et al. (2013), mostly to enlarge the sampling of a few genera. The species *Matayba tenax* was excluded, because it does not match any *Matayba* species or synonym in the Plants of the World database (https://powo.science.kew.org/taxon/urn:lsid:ipni.org:names:30007598-2), this particular terminal float in a previous analysis (Buerki et al. 2021), and the genus *Matayba* did not appear as monophyletic in previous studies (Buerki et al. 2011, 2013). The taxonomy was updated from the Plants of the World database (https://powo.science.kew.org/taxon/urn:lsid:ipni.org:names:30000506-2; PoWO 2023), replacing synonym names with accepted names. Terminals with a defined genus name but an unidentified species are replaced with sampled species of the same genus from previous studies (Buerki et al. 2011, 2013; Chery et al. 2019). The total number of terminals in the tree was 146, with the root node age of 104.37 Ma. The used tree is available as supplementary data and illustrated in the supplementary figure 2.

### Geographic data

All georeferenced preserved specimens for Sapindaceae in GBIF were downloaded (https://www.gbif.org/occurrence/download/0000527-230828120925497; GBIF.org 2023), for a total of 387,463 records. Then the taxonomy of the terminal species of the phylogenetic tree, updated with Plants of the World (PoWO 2023), was matched with the taxonomy in GBIF. As Plants of the World includes the countries in which each species was reported, the matched taxonomy is also used to keep only records sampled in the countries in which the species is native. This cleaning procedure is quite simple and does not remove all wrong records, in particular in large countries (as the filtering is done at the country level), and rejects potentially correct data in countries in which a species is not currently reported. While more careful filtering will be preferable, it provides a good enough dataset for the illustrative purpose of this example. After the filtering, 68,307 records are kept in the database. Note that this does not mean that most data is wrong in GBIF; rather, it shows that a large part of the diversity of the family was not sampled in the phylogenetic tree. As there are no geo-referenced specimen records for *Euchorium cubense*, I use a record based on a material citation for this taxon (https://www.gbif.org/occurrence/4135894102). The resulting dataset is available at https://github.com/js-arias/sapindaceae.

### Biogeographic analysis

To make inferences about the root of the tree, a root branch (Landis et al. 2013) with a length equal to 10% of the root age (i.e., 10 My) was added. Reconstructions for this root branch are ignored.

A Bayesian analysis of the λ parameter using a uniform prior is approximated in the following form: (1) The maximum likelihood value of λ was estimated; (2) a numerical integration was used to find the posterior distribution of λ using an uniform prior; (3) the posterior distribution of λ was approximated with a Gamma distribution, and 1000 random samples were taken from this Gamma distribution. In each sample, a likelihood reconstruction is performed with 100 stochastic maps, for a total of 100,000 probable histories from the posterior distribution.

To build the ancestral range maps, all the samples were smoothed using a KDE based on a spherical normal with λ = 1000 and using only the pixels in 95% of the CDF. For a better visualization, I also provide the reconstructions rotated to present day locations using the plate motion model. Lineage richness is calculated using the overlap of the reconstructions of all lineages that cross the most recent limit of a given time stage. For the speed analysis, the speed of a particular history is calculated by using the sum of the distance between the starting and ending pixels in each branch segment, measured in kilometers, and then dividing it by the length of the branch, measured in million years.

## Results

The maximum likelihood value for λ was 35.9 (logLike = -2197.005; 95% credibility interval 31.0-41.0), equivalent to a standard deviation of 1495.6 Km/My and the posterior distribution was approximated with a Gamma function with α = 108.0 and β = 3.0 (supp. Fig. 3; supp. data). The posterior average speed of a particle in the tree was 586.79 Km/My (95% CI 539.12-643.13).

Figures 2 and 3 show the main biogeographic results (see also supp. Figs. 4; reconstructions of all nodes are provided as image files in the supplementary data). In the following discussion, I use the 50% credibility interval to define the ancestral areas as well as the routes and times of dispersal (red to green interval in Figure 3). As in previous research (Buerki et al. 2011, 2013) the origin of Sapindaceae is inferred from what is today the eastern part of Asia, at the east of the Turgai Sea, at the end of the Early Cretaceous, (Fig. 2; supp. Fig. 4), but the credibility interval of 95% includes all of Eurasia and the northeastern part of North America (Fig. 2). The ancestral location inferred for all subfamilies and most tribes (except most tribes in the Haplocoeleae + Paullinieae group and Stadmanieae; supp. Fig. 4) is also the eastern part of Asia. Then, for most of its history, the main lineages of the group have remained mostly in eastern Asia and the northern hemisphere and reached tropical latitudes in relatively recent times (Figs. 2-3; supp. tables 1-6). This result is superficially similar to early results using predefined areas (Buerki et al. 2011, 2013), although the results found here have more geographic precision, both because the predefined area analyses used previously had large surfaces, and the result of their analysis produced several ambiguous assignments in the ancestral nodes. A difference from that previous research (Buerki et al. 2011, 2013) is that instead of a network of interoceanic dispersal between tropical landmasses, here the most probable solution involves multiple independent invasions of the tropics from landmasses just as north of them through land connections (Figs. 2-3; supp. tables 1- 6).

**Figure 2.**
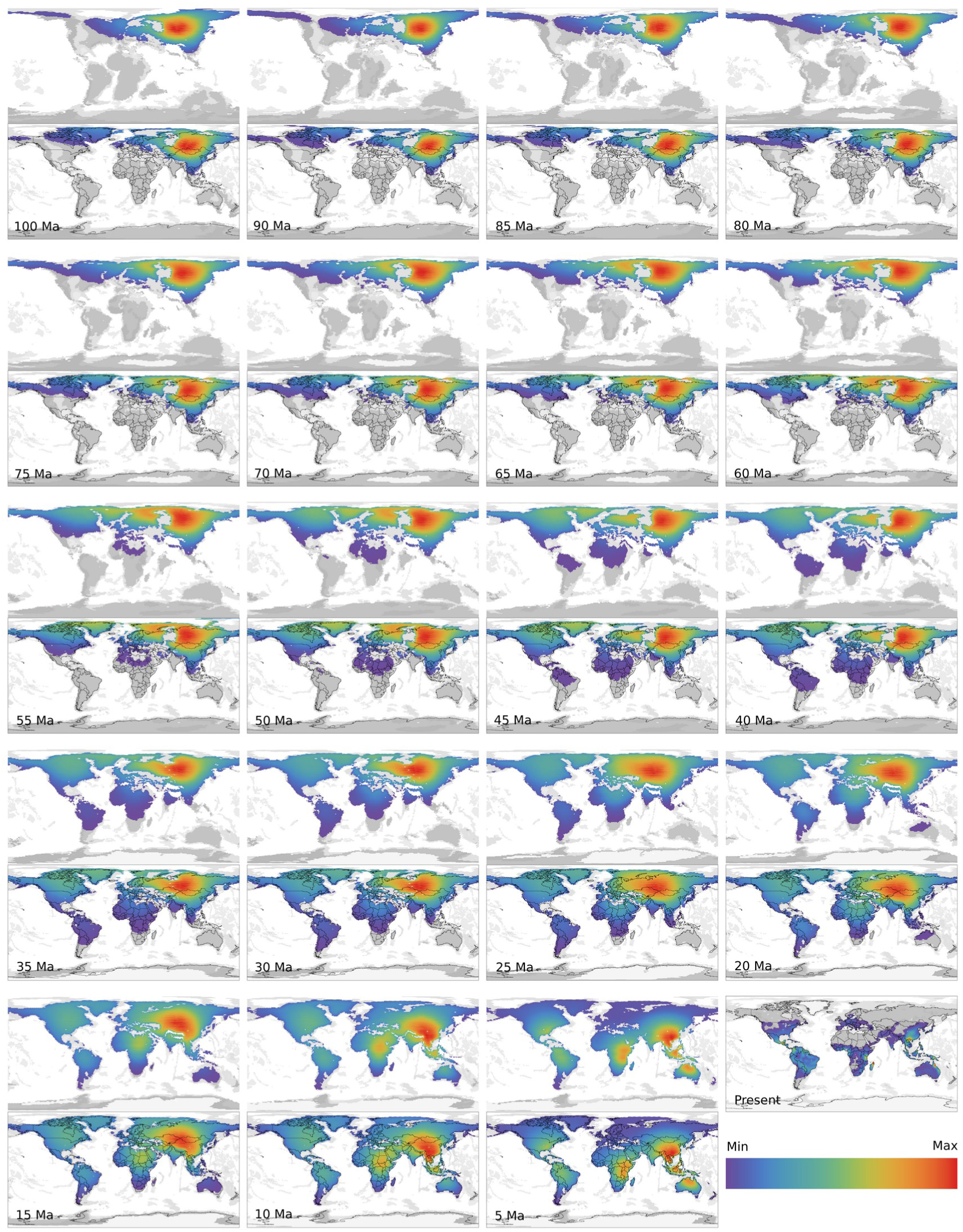
Distribution of Sapindaceae over time. The maps approximate the lineage richness by adding the posterior CDF values for all lineages alive at the end of a given time stage and scaling to the maximum value in each time stage.

**Figure 3.**
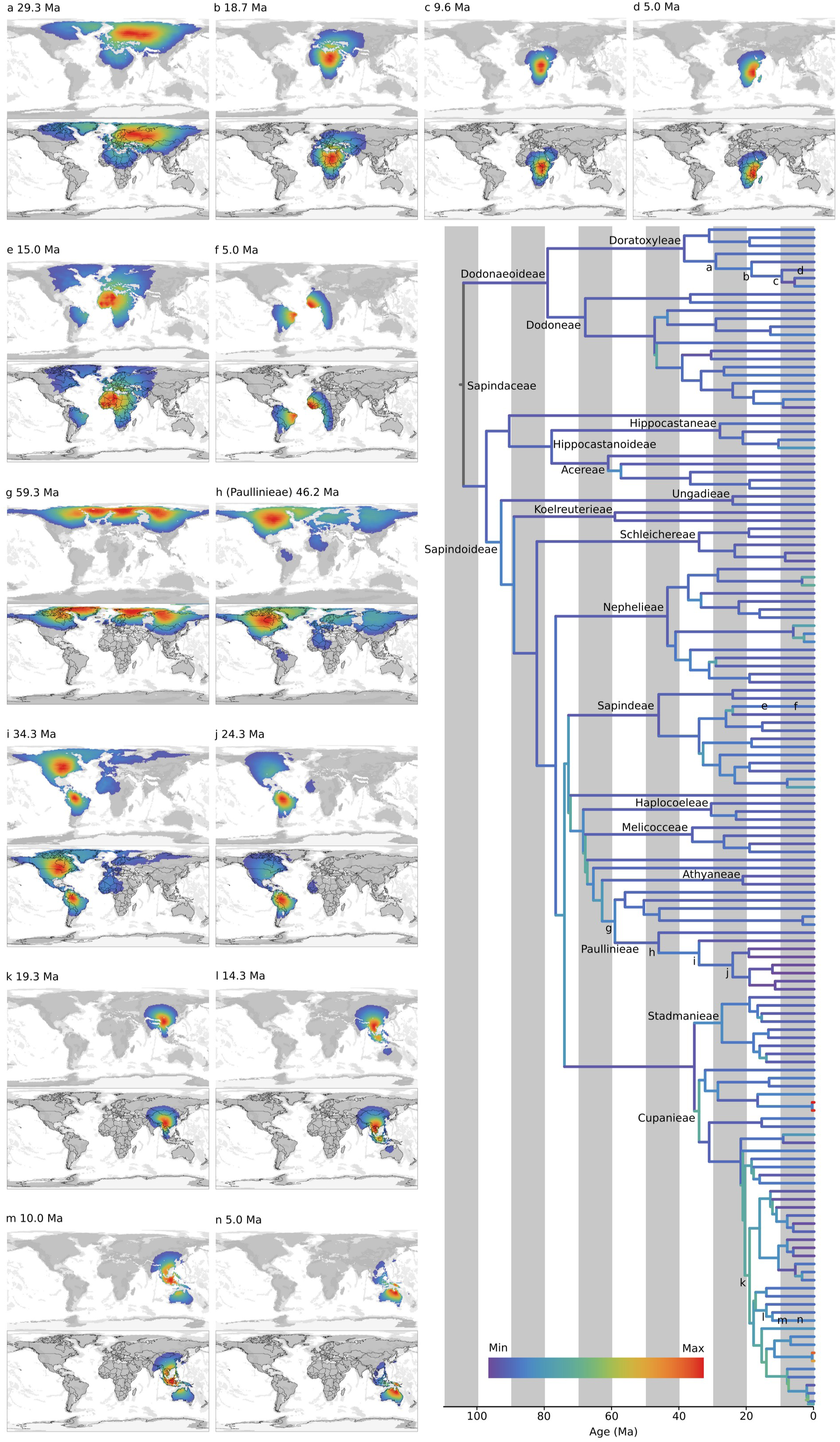
Biogeographic history of Sapindaceae. Sapindaceae phylogenetic tree with branches shaded by their posterior mean velocity in Km/My and log10 transformed. The maps show reconstructions of 95% of the cumulative density function (CDF) of the posterior at particular nodes, with shading reflecting the CDF value under a paleogeographic model (top) and projected in the current geography (bottom).

As in previous research (Buerki et al. 2011, 2013), an early dispersal involves a trip from Eurasia to North America, but the date of these initial dispersals is most recent: it is in the 145-80 Ma time slice in previous research (Buerki et al. 2011, 2013). Here, the earliest dispersal begins at 68 Ma and continues until very recent times (supp. table 1). The most probable land route for these dispersals is through Europe and Greenland, using the De Geer land route (Fig. 3g-h; supp. table 1). Many of these dispersals use North America as an intermediate stop on trips that end in Central America, the Caribbean, or South America (Fig. 3g-i). Most lineages in South America originate from North American ancestors (supp. table 2). They reach South America through two routes: an ancient route (45-20 Ma) that uses ancient Central America (Fig. 3g-j; supp. table 2), and a more recent one using either Central America or the Caribbean (15-0 Ma; supp. table 2). It is inferred in a few lineages that an Atlantic crossing from Africa occurred in recent times (15-0 Ma, Fig. 3e-f; supp. table 2), although there is some ambiguity as a proposed alternative is a dispersal using the De Geer pass. Different from previous results (Buerki et al. 2011, 2013), there is no instance of a dispersal from Australia to South America, either through Antarctica or using the West Wind Drift.

There are some early dispersals into Africa (50-35 Ma; supp. table 3), centered around what is today the Mediterranean. Most of the dispersals into Africa start at the Oligocene (30-20 Ma), and the main route is using Anatolia peninsula and what today is Iran through the Arabian peninsula to reach Africa (fig. 3a-b; supp. table 3). The majority of the lineages in Madagascar originated from an African ancestor and reached Madagascar in quite recent times (10-0 Ma; supp. table 4). Contrary to previous research (Buerki et al. 2011, 2013) that proposes a large amount of dispersal from Madagascar into Africa, no instance was inferred here. Also, there was no support for a route connecting Africa and Southeast Asia via Madagascar and India during the Paleocene-Eocene. Two recent dispersals involving very short branches with sister groups far away indicate a potential Indian ocean dispersal, one to Madagascar (sister in New Guinea) and the other in West Africa (sister in New Caledonia). As these branches are small (less than a million year) and their descendants are separated by the ocean, the ancestors are inferred from the Malesia archipelago and even Kerguelen, and the speed of these branches is far greater than any other branch (Fig. 3), from 5300 to 13200 Km/My (supp. data).

While previous work set the first dispersal into Australia in the Paleocene to Eocene time slice (61-33 Ma; Buerki et al. 2011, 2013), here the oldest movement into what is today Australia and New Guinea was inferred in the Miocene (20 Ma; supp. table 5). All of the dispersals are through the Malesia archipelago (Fig. 3k-n). Contrary to previous research (Buerki et al. 2011, 2013), no dispersal from Australia into Asia was found. The origin of the New Caledonian taxa is left out in previous research as it is inside a predefined area with Australia (Buerki et al. 2011, 2013). Here, the origin of New Caledonian taxa is quite recent (<5 Ma), and from Australia or New Guinea (Fig. 3n; supp. table 6), the only exception is the already mentioned small branch connecting a New Caledonian taxon with a West African taxon.

The fastest branches are found in the Cupanieae tribe, including the two faster groups of the Indian Ocean disjunction (Fig. 3). While not as fast, many of the basal branches of this tribe reached relatively high speeds (1000-1500 Km/My), which is consistent with their recent time of origination (<30 Ma) and their widespread presence in Africa, Madagascar, Australia, and South America. While previous research set the ancestral area of the group in Australia, and then a lot of dispersal using the West Wind Drift (Buerki et al. 2011, 2013), the history found here started in Asia, and then mostly land routes were used to quickly reach their current distribution (Fig. 3k-n).

## DISCUSSION

### Plate Tectonics

While paleogeography and plate tectonics have been fundamental to phylogenetic biogeography since its inception (Brundin 1965, 1966, 1972; Nelson 1969, 1974), it was not until the implementation of the DEC model (Ree et al. 2005; Ree and Smith 2008; Webb and Ree 2012; Landis 2016) that these concepts were fully integrated into the inference machinery of phylogenetic biogeography. Unfortunately, DEC was developed under the paradigm of predefined areas, and therefore its paleogeographic models inherit the problems of these methods, including poor spatial resolution, but add some more (Ree and Sanmartín 2009). As there are few areas, the temporal resolution of the models is also poor; for example, the most complex predefined area model (Landis 2016) uses only 25 areas for the whole world and 26 time slices for the whole Phanerozoic (540 Myr, i.e., on average one time slice every 20 Myr). Given their low resolution, predefined areas mask the complex history of the tectonic elements that make up these predefined areas; a paradigmatic example is the “Oriental region” in southeast Asia in most global models (e.g., Buerki et al. 2011, 2013; Webb and Ree 2012; Landis 2016; Kawahara et al. 2023). Usually users are responsible for building the dispersal matrix, which is cumbersome for models with a large number of predefined areas and requires subjective and poorly documented choices that can be critical to final results, such as defining how well connected two areas are.

Following previous authors (Ree et al. 2005; Ree and Smith 2008; Webb and Ree 2012; Landis 2016), this paper advocates for the integration of paleogeography models as the best way to incorporate the Earth’s deep time history into the analysis. However, I take a step further and propose that these models should be the same models used by geologists for their paleogeographic reconstructions (e.g., Merdith et al. 2021; Müller et al. 2019, 2022) under an explicit geographic data model. As a way of contrast with predefined area models, the paleogeographic model used in the empirical part of this paper (Cao et al. 2017; Müller et al. 2022) is defined for 109 time slices for the Phanerozoic (each time slice has a duration of 5 million years) and about 18,500 pixels in each time slice, which represent an increase of four times in the temporal scale and three orders of magnitude in the spatial scale relative to the most detailed predefined area model (Landis 2016).

As O’Donovan et al. (2018) did not use a paleogeographic model in their analysis of dinosaurs, they suggested that the effect of plate motion is minimal because it is a global group and biological and ecological forces are more influential than plate movement. However, plate tectonics affects not only the speed and direction of the movement of taxa but also the opportunities and limitations of taxon movement due to changes in spatial configuration. As such, it underlies biological and ecological forces that shape the taxon’s movement. O’Donovan et al. (2018) also argue that directional movement implies that most movement is biological. But the only way to separate tectonic movement from biological movement is by including both types of movement in the model. In the model presented here, tectonic movement (provided by the plate motion model) is explicitly separated from biological movement (inferred using the diffusion model).

The use of paleogeography models in biogeography is sometimes criticized for giving less weight to biogeographic evidence relative to geological evidence (e.g., Parenti and Ebach 2009). They argue that biogeographic analysis should be used to propose new paleogeographic hypotheses. As far as I know, there are few explicit attempt to transform the results of a biogeographic analysis into a plate motion model (e.g., Meert and Lieberman 2004; Bolotov et al. 2022), but they are based on a small datasets, and the methodology to move from a biogeographic hypothesis to a paleogeographic model is not well defined. A more achievable goal is to compare the likelihood of two or more paleogeography models using many clades (as proposed by Ree and Smith 2008; Webb and Ree 2012). However, such comparisons require the same plate motion model used by geologists (as in the method proposed here) in order to accurately test specific paleogeographic reconstructions.

A potential limitation of using time slices is that it increases the required computational resources (Bielejec et al. 2014; Landis 2016). These are because conditional likelihoods need to be calculated for each internode and values for each pixel need to be stored in memory. The model used in the empirical analysis has relatively close time stages (each 5 Myr), and while closer time stages will be preferable, it seems that it is good enough to provide reasonable answers to a typical biogeographic problem and to be treatable with current computer power.

### Spherical Earth

Early attempts to incorporate a spherical Earth model into phylogenetic biogeography often fell short due to various limitations. Some studies used great circle distances without scaling the probability with the area of available destinations in a sphere (Lemmon and Lemmon 2008; Lemey et al. 2010; Pybus et al. 2012; Landis et al. 2013), while others made a 3D representation of the Earth but failed to use great circle distances in the likelihood calculations (O’Donovan et al. 2018). While the difference with a full spherical model will be minimal for cases where the λ parameter is large, either because the lineages have a slow diffusion or the branch lengths are small, or when the geographic span of a clade is quite restricted (as could be the case of some intra-specific phylogeographic analyses), they may not be appropriate for inter-specific phylogenetic biogeography analyses with long time scales, many long branches, and widespread distributions (this last case is also possible in intra-specific phylogeography). In such scenarios, it is critical to use a spherical diffusion approach (Bouckaert 2016; Louca 2021), even if it sometimes requires more complex computations.

Using a flat Earth model can lead to some unintended consequences, such as applying methods that work well in flat Euclidean geometry but not on a spherical Earth. For example, some authors use latitude as a trait value, in which a particle has the same probability to move to the north or south (e.g., Silvestro et al. 2019), but because of Earth’s sphericity, this probability is dependent on latitude. Another common scenario is to model the movement in latitude and longitude as diffusion on independent axes (Lemey et al. 2010; Pybus et al. 2012; Quintero et al. 2015) or compare the bearing from points in different locations (Lemmon and Lemmon 2008; O’Donovan et al. 2018). On flat Euclidean geometry, this can be done, as these measures are made relative to coordinate axes, so, for example, rotating the axes will rotate all relationships in the same magnitude. On Earth, latitude and bearing are measured relative to a particular point (usually the North Pole), and any change in this point will produce changes in coordinates and bearings of different magnitudes. This is the same reason that makes it impossible to compare the covariance matrices of the anisotropic spherical normal at different points (Hauberg 2018).

### Landscape

Although the model presented here utilizes some landscape information (pixel final positions are conditioned by a prior in the pixel), it is not a full landscape model, as particles have unrestricted movement in any pixel. A better alternative is to condition particle movement according to the type of pixel it traverses. This landscape model has been proposed for phylogeographic analysis (Bouckaert et al. 2012). However, it incurs a significant computational cost, with a significant reduction in the geographic resolution of the model.

Despite its limitations, the landscape model used here has some influence on the inference, as shown in the empirical example, where some dispersal routes can be inferred above emerging land instead of a more “direct” path over the ocean. As the values of λ become larger, the movement slows down; therefore, there will be more influence from the landscape as movement will be restricted to nearly suitable pixels. In fact, the Bouckaert et al. (2012) method is based on the numerical integration of small time steps and very slow diffusion (analogous to a large λ value). This observation is consistent with what we expect: the higher the capability of dispersal, the fewer the restrictions on movement because of geographic barriers. There is another consequence of the use of a landscape: as pixels near the boundaries have fewer suitable pixels, the ancestral pixels are most probably assigned far from the boundaries, which is more noticeable when the ancestral area is on a large landmass.

From a computational standpoint, a model without landscape allows for faster computation, either by using contrasts (but losing ancestral locations Louca 2021), only storing the means (Bouckaert 2016), traditional MCMC with data augmentation (Bouckaert 2016; O’Donovan et al. 2018), or heuristic searches of the maximum likelihood (Lemmon and Lemmon 2008). But as the effects of the landscape are not linear, they cannot be solved analytically, which requires full integration over all pixels. Fortunately, at least for the simple landscape model used here and shown with the empirical example, current computational power is generally adequate for most common cases and provides a good approximation of the effect of landscape connectivity.

### Diffusion

The diffusion model has been criticized because uncertainty in assignment increases with time (Ronquist and Sanmartín 2011; Quintero et al. 2015). For fast-moving groups, it will be difficult to find a meaningful answer, but this problem affects any other biogeographic method. Previous uses of the diffusion model (Bouckaert et al. 2012; Bouckaert 2016; Nylinder et al. 2016; O’Donovan et al. 2018; Swenson et al. 2019), as well as the empirical example presented in this paper, show that it is possible to use the diffusion model on both large temporal and geographic scales.

Another criticism of the diffusion model is that it is unsuitable for taxa living on islands (Lemmon and Lemmon 2008; Quintero et al. 2015). This criticism arises from two factors: first, the diffusion process is modeled as homogeneous across the Earth surface, and second, it fails to account for founder event speciation. However, as shown in the empirical example and discussed in the landscape section before, the coupling of a diffusion model with a landscape model (even if the landscape model is not complete) can provide reasonable results in the presence of a heterogeneous landscape (see also Swenson et al. 2019). The current diffusion model does not account for founder event speciation. This event remains a topic of controversy even in predefined area models (Ree and Smith 2008; Ree and Sanmartín 2018; Matzke 2022; Landis et al. 2022). But as seen in the empirical example, the diffusion model was able to detect the movement speedup that is usually associated with founder events.

Johansson et al. (2018) criticized the diffusion model for its inability to assess the strength of the biogeographical signal in its comparison with DEC. However, similar to predefined area methods or any likelihood-based analysis, the signal strength in a diffusion model is determined by the size of the interval of the assignments. If there are several peaks in the posterior or the pixels with the highest posterior spread over a large surface, it indicates a weak signal. Conversely, if the most pixels cluster around a small posterior peak, it indicates a strong signal.

### Explicit Geography

Methods based on explicit geographic ranges use all available geographic information in the terminal data. As already discussed in the literature, there are several advantages over methods based on predefined areas (Arias et al. 2011; Landis et al. 2013; Quintero et al. 2015; Arias 2017; O’Donovan et al. 2018).

Explicit geography methods allow for detailed geographic assignments both as split nodes and along specific parts of a lineage, providing a more detailed view of the evolution of the geographic range of each lineage of the studied phylogeny (Lemmon and Lemmon 2008; Arias et al. 2011; Bouckaert et al. 2012; Pybus et al. 2012; Landis et al. 2013; Quintero et al. 2015; Bouckaert 2016; Arias 2017; Johansson et al. 2018; O’Donovan et al. 2018). This explicit geographic assignment offers the opportunity to test more detailed hypotheses about the space and evolution of lineages. Here are three examples, that can be derived from a comparison of the empirical example with previous analysis of the same group using predefined area methods (Buerki et al. 2011, 2013). (*i*) With explicit geography methods, specific geographic locations are inferred as the ancestral location at each node, and even when the assignment is ambiguous (i.e., a large surface), its geographic scope is clearly delimited. As predefined areas used in methods such as DEC are usually large surfaces, their users must attempt ad hoc procedures to improve the geographic scale of their analyses (e.g., Smith and Donoghue 2010). (*ii*) In predefined area methods, such as DEC, the potential connection between areas is defined either a priori or left free, without any geographic constraint (Ree et al. 2005; Ree and Smith 2008), which discards geographic knowledge. As shown in the empirical example, with methods based on diffusion, it is possible to infer the potential dispersal pathway connecting the ancestor and descendant points. These paths are detected from the data and not defined a priori, and explicitly include the constraints of the geography. (*iii*) Spatially explicit models can infer distances and speeds from the movement through the phylogeny; this is impossible to measure with predefined area methods, as they discard any movement inside a predefined area (Ronquist and Sanmartín 2011) and resort to ad hoc methods to measure the distance between two predefined areas (Landis et al. 2022).

Ronquist and Sanmartín (2011) argue that methods based on explicit geography treat all changes as equal, then common movement within an area can be “saturated” (as molecular data), and “swamp” the rare and more informative movement between areas. This characterization seems to be based on a confusion of unconstrained predefined area models (which treat all changes as equal Ronquist 1997; Ree et al. 2005; Ree and Smith 2008) with a diffusion model, in which movements are constrained by the distance between points. As moving to farther distances is more unlikely, any unexpected large-distance movement required by the data will be quickly detected as an increase in diffusion speed (fig. 3). On the other hand, if the movement is between close areas, the diffusion movement will infer this movement as simple diffusion (see the empirical example).

To make use of explicit geographic data, high-quality data is required for the terminal distributions. Fortunately, centuries of careful taxonomic work, as well as modern open-access databases and repositories such as GBIF (https://www.gbif.org/) and FossilWorks (http://www.fossilworks.org/), provide a large amount of digitalized spatial data associated with specific taxa. However, these datasets often require cleaning and are not always complete. Despite the potential for error and incomplete geographic distribution data, using large, predefined areas is not a viable solution. Doing so would discard the information content of these scarce but valuable data points (Landis et al. 2013). Even if a researcher is hesitant to give weight to a few observations, it is possible with the current method (as in previous ones Bouckaert et al. 2012; Nylinder et al. 2014; Bouckaert 2016; Louca 2021) to assign a large geographic area to a terminal around the observed point(s), thus reducing the weight of the data. This approach differs from using predefined areas in two critical ways: it takes geographic locations into account, and the assigned area can be different for each terminal.

Similar to the method presented by Quintero et al. (2015), the current implementation avoids Monte Carlo integration of tip data. However, the present approach goes further by avoiding Monte Carlo integration of augmented data at internal nodes as well. This step does make calculations slower compared to a single iteration with data augmentation. Nonetheless, the large number of iterations required by the Monte Carlo integration of the augmented node data to produce a good sample of the pixel likelihoods at each node reduces the benefits of this acceleration. In particular, incorporating paleogeography models requires estimation in many intermediate branches, which can hamper the convergence of the Monte Carlo integration of augmented data (Pybus et al. 2012).

### Limitations

In the previous sections, I highlighted some of the limitations of the method presented here, including the fact that landscape information is not used in its entirety. However, there are other limitations to the method that are worth mentioning.

The current implementation employs a strict clock model, with a single diffusion rate used throughout the entire tree. This limitation is shared with most biogeographic methods based on statistical models (Ree et al. 2005; Lemmon and Lemmon 2008; Ree and Smith 2008; Landis et al. 2013; Quintero et al. 2015; Landis 2016), and even with this limitation, they are used for empirical research on large and ancient clades (e.g., Buerki et al. 2011, 2013; Dupin et al. 2017; Landis et al. 2021; Bolotov et al. 2022; Kawahara et al. 2023). The main issue with a strict clock is that it can distort the inference when fast and slow-moving lineages are mixed. As demonstrated in the empirical example, many branches move at different speeds, so it will be fruitful to implement a relaxed clock procedure (e.g., Lemey et al. 2010; O’Donovan et al. 2018) in the future.

The method presented here does not estimate or use widespread ancestral ranges, which prevents the inference of cladogenetic events and the inference of the extinction process. The method implicitly assumes sympatric speciation. While potential allopatric scenarios might be detected by examining the results closely, it would be preferable if the model included these events in its inference machinery. While estimating extinction is desirable, it is worth noting that this is a challenging parameter to estimate in most biogeographic methods (e.g., Ronquist 1997; Ree et al. 2005; Ree and Smith 2008; Matzke 2022), and how to couple it with the diffusion model remains an open question.

While producing maps with ancestral ranges is more satisfactory than the abstract results of methods based on predefined areas, they are more challenging to display, as usually they require a full map (and as the geography has changed a lot, results rotated to present locations are required to understand the geography). Note that this is also a problem in predefined areas when the number of area combinations is large and it is difficult to differentiate between the colors of the different reconstructions (the usual result presented in those analyses). In a similar way, if age uncertainty is to be taken into account, it is not clear how to represent the results when the age ranges of a node cross more than one time stage, as the assigned pixels will only have meaning in a particular time stage.

## CONCLUDING REMARKS

The method presented in this paper is the first to incorporate an explicit plate motion model into an explicitly geographic phylogenetic biogeography method. The method also uses a spherical Earth (previously used by Bouckaert 2016; O’Donovan et al. 2018; Louca 2021) and considers landscape information (although it is not a full landscape implementation; Bouckaert et al. 2012) that changes with time. The method is flexible and can use data from sample locations or range maps, both homogeneous (like a distribution map) and heterogeneous (like a distribution model). It can also be used for intra-specific phylogeography as well as deep-time phylogenetic biogeography.

The method presented here provides several possible avenues of inference not explored in this paper. For example, if the reconstructions are not of interest, it may be possible to optimize multiple trees from a posterior set with a low-resolution model, say an e120, to provide a biogeographic dating estimation (Webb and Ree 2012; Landis 2016). If there are multiple phylogenies, it is possible to study changes in the richness gradient in a similar way to that illustrated in Figure 2.

Explicit geography methods are an improvement over predefined area methods because they take into account the data from the distribution range of the terminals and use high-resolution models of the spherical Earth, which make better use of biological and paleogeographic data to explain the evolutionary history of the geographic ranges.

## DATA AVAILABILITY STATEMENT

Supplementary material and data are available on a GitHub repository https://github.com/js-arias/sapindaceae for the main data and raw results. The code of the *PhyGeo* implementation is available at https://github.com/js-arias/phygeo.

## ACKNOWLEDGEMENTS

I have received a lot of support from J. Grosso, P. Goloboff, C. Szumik, A. Galvis, S. Catalano, D. Casagranda, J. Hyvönen, L. Amador, P. Giannini, S. Bertelli, J. Fratani, J. Flores, and J. Daza, who were critical of encouraging me to finish this work, or read multiple version of the manuscript. G. Dantur was the first user of *PhyGeo*. Preliminar versions of this project have been presented in several scientific meetings in Argentina, Colombia, Ecuador, Finland, and U.S.A., and I am grateful for all the colleagues in these meetings who take the time to show me their interest, give me suggestions, and ask me questions about the method.

## FUNDING

This work was supported by the Ministerio de Ciencia y Técnica de Argentina (PICT 2020-02650).

## Notes

https://github.com/js-arias/phygeo/

https://github.com/js-arias/geomodels

https://github.com/js-arias/sapindaceae

### Competing Interest Statement

The authors have declared no competing interest.

