## Supplementary material for "Phylogenetic Biogeography Inference Using Dynamic Paleogeography Models and Explicit Geographic Ranges": supp. data

J. Salvador Arias[1,2,*]

[1]*Unidad Ejecutora Lillo (CONICET-Fundación Miguel Lillo), S.M. de Tucumán, Tucumán, Argentina*

[2]*Current address: Laboratorio de Genética Evolutiva, Instituto de Biología Subtropical (CONICET-UnaM), Universidad Nacional de Misiones, Felix de Azara 1552, CP 3300, Posadas, Misiones, Argentina*

### Supplementary Information

**Supplementary table 1.** Dispersals into North America. The node column indicates the node that contains the youngest part of the credibility interval (of 50%); refer to supplementary figure 3 to get the node IDs. The time frame column reports an approximation of the 50% credibility interval for the dispersal (i.e., the oldest to youngest dispersal instances). Events indicated as ambiguous indicate that there are alternative dispersal routes for these nodes.

| Node | Time Frame 50% CI (Ma) | Source | Comments |
|---|---|---|---|
| 149 | 68.7 – 15.0 | Eurasia | De Geer pass |
| 18 | 68.1 – 15.0 | Eurasia | De Geer pass |
| 30 | 40.0 – 15.0 | Eurasia | De Geer pass |
| 129 | 33.1 – 20.0 | Eurasia | De Geer pass |
| 4 | 30.0 – 10.0 | Eurasia | De Geer pass |
| 63 | 30.0 – 5.0 | Eurasia | De Geer pass |
| 9 | 29.3 – 10.0 | Eurasia | De Geer pass |
| 51 | 28.1 – 10.0 | Eurasia | De Geer pass |
| 69 | 24.3 – 10.0 | Eurasia | De Geer pass, or Bering pass |
| 138 | 15.0 – 0.0 | Eurasia | Bering pass, or De Geer pass |
| 273 | 15.0 – 5.0 | Eurasia | De Geer pass (ambiguous) |
| 282 | 14.3 – 10.0 | Eurasia | De Geer pass (ambiguous) |
| 56 | 10.0 – 5.0 | Eurasia | De Geer pass |

**Supplementary table 2.** Dispersals into South America. See supp. table 1 for an explanation of the columns.

| Node | Time frame 50% CI (Ma) | Source | Comments |
|---|---|---|---|
| 162 | 40.0 - 21.1 | North America | Throu Central America (ambiguous) |
| 177 | 40.0 - 10.0 | North America | Throu Central America |
| 160 | 30.0 - 5.0 | North America | Throu Central America (ambiguous) |
| 167 | 30.0 - 5.0 | North America | Throu Central America |
| 20 | 25.0 - 5.0 | North America | Throu Central America and Caribbean (ambiguous) |
| 154 | 25.0 - 0.0 | North America | Throu Central America and Caribbean |
| 282 | 14.3 - 8.1 | Africa | Throu Atlantic ocean (ambiguous) |
| 20 | 15.0 - 0.0 | Africa | Throu Atlantic ocean (ambiguous) |
| 30 | 15.0 - 0.0 | North America | Throu Central America and Caribbean |
| 123 | 15.0 - 0.0 | Africa | Throu Atlantic ocean |
| 131 | 15.0 - 0.0 | North America | Throu Central America and Caribbean |
| 151 | 15.0 - 0.0 | North America | Throu Central America |

| | | | |
|---|---|---|---|
| | | | and Caribbean |
| 153 | 15.0 - 0.0 | North America | Throu Central America and Caribbean |
| 171 | 15.0 - 0.0 | North America | Throu Central America and Caribbean |
| 176 | 10.0 - 0.0 | North America | Throu Central America and Caribbean |
| 273 | 10.0 - 0.0 | North America or Africa | Throu Atlantic ocean, or Caribbean |
| 283 | 8.1 - 5.0 | North America | Throu Central America and Caribbean (ambiguous) |
| 286 | 7.7 - 2.3 | North America | Throu Central America and Caribbean (ambiguous) |

**Supplementary table 3.** Dispersals into Africa. See supp. table 1 for an explanation of the columns.

| Node | Time frame 50% CI (Ma) | Source | Comments |
|---|---|---|---|
| 144 | 50.0 - 30.6 | Eurasia | Throu eastern Mediterranean (ambiguous) |
| 172 | 35.0 - 10.0 | Eurasia | Throu wastern Mediterranean, or eastern Mediterranean |
| 191 | 35.0 - 18.1 | Eurasia | Throu Anatolia, Iran, and Arabian peninsula |
| 107 | 35.0 - 5.0 | Eurasia | Throu eastern Mediterranean |
| 121 | 34.3 - 5.0 | Eurasia | Throu eastern Mediterranean |
| 106 | 30.0 - 5.0 | Eurasia | Throu Anatolia, Iran, and Arabian peninsula |
| 145 | 30.0 - 5.0 | Eurasia | Throu western Mediterranean (ambiguous) |
| 146 | 30.0 - 5.0 | Eurasia | Throu eastern Mediterranean (ambiguous) |
| 10 | 29.3 - 15.0 | Eurasia | Throu eastern Mediterranean (ambiguous) |
| 20 | 25.0 - 10.0 | Eurasia | Throu western Mediterranean (ambiguous) |
| 25 | 25.0 - 5.0 | Eurasia | Throu Anatolia, Iran, and Arabian peninsula |
| 133 | 25.0 - 5.0 | Eurasia | Throu Anatolia, Iran, and Arabian peninsula (ambiguous) |
| 23 | 25.0 - 0.0 | Eurasia | Throu Anatolia, Iran, and Arabian peninsula |
| 100 | 20.0 - 3.1 | Eurasia | Throu Anatolia and Arabian peninsula |
| 26 | 20.0 - 0.0 | Eurasia | Throu Anatolia, Iran, and Arabian peninsula |
| 89 | 20.0 - 0.0 | Eurasia | Throu Anatolia and Arabian peninsula |
| 7 | 19.3 - 5.0 | Eurasia | Throu Anatolia, Iran, and Arabian peninsula |
| 272 | 18.1 - 6.9 | Eurasia | Throu Anatolia, Iran, and Arabian peninsula |

| | | | |
|---|---|---|---|
| 218 | 0.6 - 0.0 | Kerguelen or Eurasia | Indian ocean |

**Supplementary table 4.** Dispersals into Madagascar. See supp. table 1 for an explanation of the columns.

| Node | Time frame 50% CI (Ma) | Source | Comments |
|---|---|---|---|
| 25 | 10.0 - 0.0 | Africa | Throu Monzambique channel |
| 193 | 10.0 - 0.0 | Africa | Throu Monzambique channel |
| 201 | 10.0 - 0.0 | Africa | Throu Monzambique channel |
| 202 | 10.0 - 0.0 | Africa | Throu Monzambique channel |
| 206 | 10.0 - 0.0 | Africa | Throu Monzambique channel |
| 207 | 10.0 - 0.0 | Africa | Throu Monzambique channel |
| 13 | 9.6 - 0.0 | Africa | Throu Monzambique channel |
| 15 | 5.0 - 0.0 | Africa | Throu Monzambique channel |
| 16 | 5.0 - 0.0 | Africa | Throu Monzambique channel |
| 28 | 5.0 - 0.0 | Africa | Throu Monzambique channel |
| 133 | 5.0 - 0.0 | Africa or Inida | Throu Monzambique |

| | | | |
|---|---|---|---|
| | | | channel, or Indian ocean |
| 173 | 5.0 - 0.0 | Africa | Throu Monzambique channel |
| 174 | 5.0 - 0.0 | Africa | Throu Monzambique channel |
| 195 | 5.0 - 0.0 | Africa | Throu Monzambique channel |
| 198 | 5.0 - 0.0 | Africa | Throu Monzambique channel |
| 204 | 5.0 - 0.0 | Africa | Throu Monzambique channel |
| 277 | 5.0 - 0.0 | Africa | Throu Monzambique channel |
| 278 | 5.0 - 0.0 | Africa | Throu Monzambique channel |
| 280 | 0.6 - 0.0 | Eurasia | Indian ocean |

**Supplementary table 5.** Dispersals into Australia and New Guinea. See supp. table 1 for an explanation of the columns.

| Node | Time frame 50% CI (Ma) | Source | Comments |
|---|---|---|---|
| 212 | 20.0 - 5.0 | Eurasia | Throu Malesia |
| 239 | 19.3 - 0.0 | Eurasia | Throu Malesia |
| 38 | 15.0 - 5.0 | Eurasia | Throu Malesia |
| 216 | 15.0 - 5.0 | Eurasia | Throu Malesia |
| 229 | 5.0 - 0.0 | Eurasia | Throu Malesia |
| 40 | 15.0 - 0.0 | Eurasia | Throu Malesia |
| 43 | 15.0 - 0.0 | Eurasia | Throu Malesia |
| 210 | 15.0 - 0.0 | Eurasia | Throu Malesia |
| 268 | 14.3 - 0.0 | Eurasia | Throu Malesia |
| 271 | 12.5 - 5.0 | Eurasia | Throu Malesia |
| 270 | 12.5 - 0.0 | Eurasia | Throu Malesia |
| 226 | 9.3 - 5.0 | Eurasia | Throu Malesia |
| 37 | 10.0 - 0.0 | Eurasia | Throu Malesia |
| 42 | 10.0 - 0.0 | Eurasia | Throu Malesia, skip New Guinea |
| 234 | 10.0 - 0.0 | Eurasia | Throu Malesia |
| 237 | 10.0 - 0.0 | Eurasia | Throu Malesia |

| | | | |
|---|---|---|---|
| 265 | 10.0 - 0.0 | Eurasia | Throu Malesia |
| 266 | 10.0 - 0.0 | Eurasia | Throu Malesia |
| 6 | 5.0 - 0.0 | Eurasia | Throu Malesia |
| 222 | 5.0 - 0.0 | Eurasia | Throu Malesia |
| 232 | 5.0 - 0.0 | Eurasia | Throu Malesia |
| 281 | 0.6 - 0.0 | Eurasia | Throu Malesia |

**Supplementary table 6.** Dispersals into New Caledonia. See supp. table 1 for an explanation of the columns.

| Node | Time frame 50% CI (Ma) | Source | Comments |
|---|---|---|---|
| 38 | 5.0 - 0.0 | Australia or New Guinea | |
| 213 | 5.0 - 0.0 | Australia or New Guinea | |
| 216 | 5.0 - 0.0 | Australia or New Guinea | |
| 229 | 5.0 - 0.0 | Australia or New Guinea | |
| 247 | 5.0 - 0.0 | Australia | |
| 258 | 5.0 - 0.0 | Australia | |
| 271 | 5.0 - 0.0 | Australia or New Guinea | |
| 260 | 3.7 - 0.0 | Australia | |
| 261 | 3.7 - 0.0 | Australia | |
| 219 | 0.6 - 0.0 | Kerguelen or Eurasia | Throu Malesia |

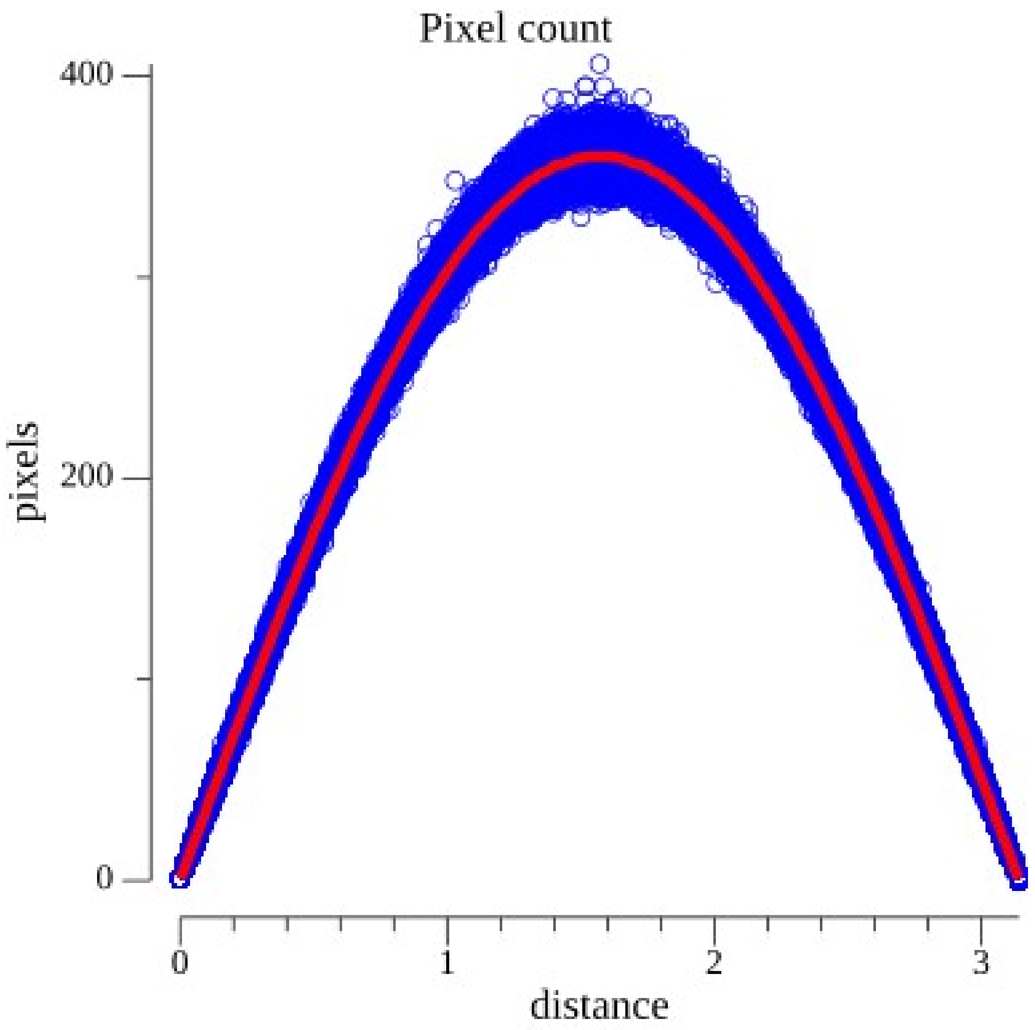


***Supplementary figure 1***. *Plot of the number of pixels at a given distance (in radians) from 20 random pixels in each ring of an e360 pixelation used in this paper; the pixelation has 180 rings (for a total of 3600 sampled pixels). In red is the number of pixels at a given distance from the North Pole, which is used for quick approximations of the spherical normal in the likelihood calculations.*

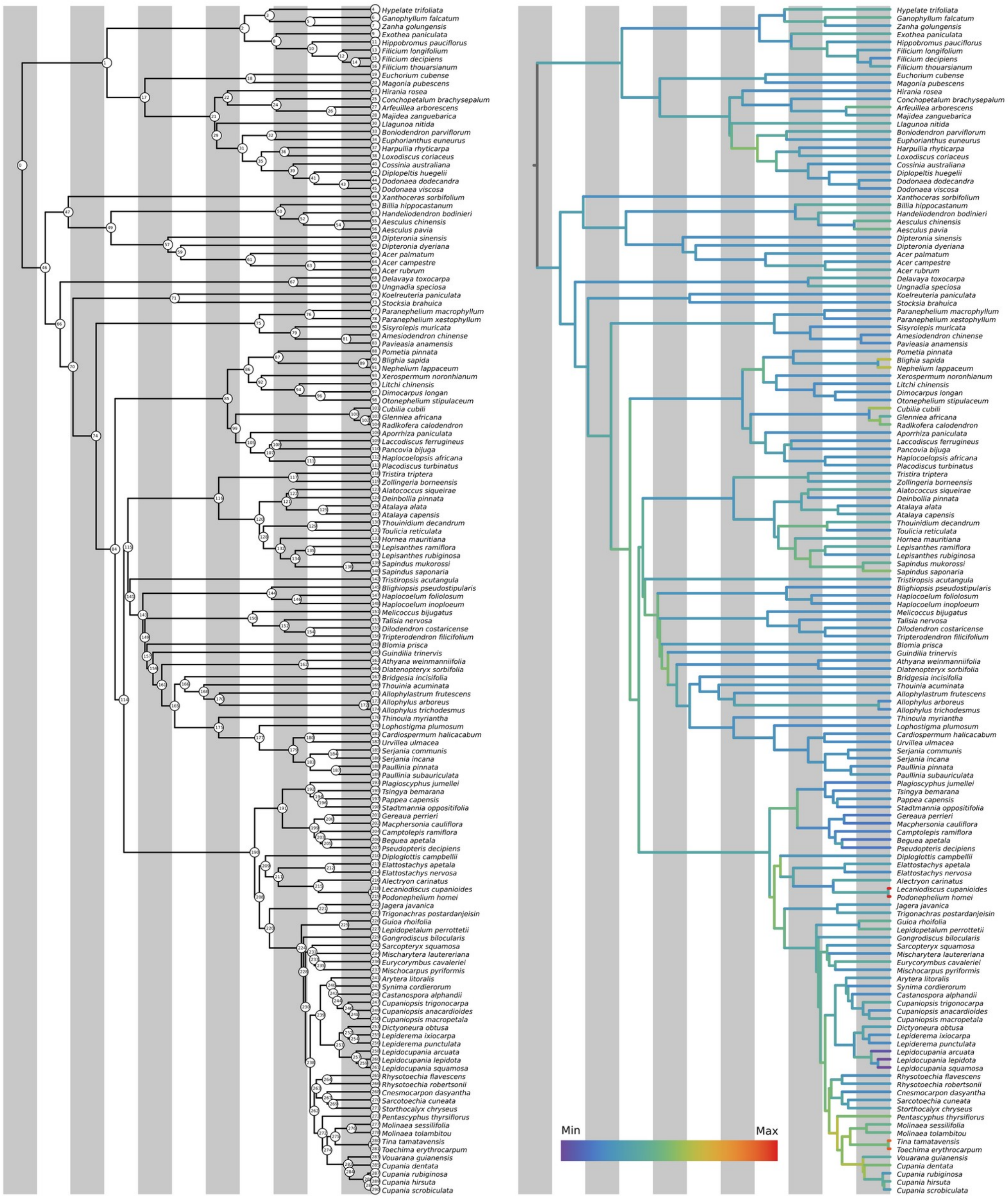


***Supplementary figure 2***. *Phylogenetic relationships of Sapindaceae used in this work. On the left, the tree with the node numbers indicated; on the right, the same tree with the branches colored by its average speed (in Km/My) log10 scaled.*

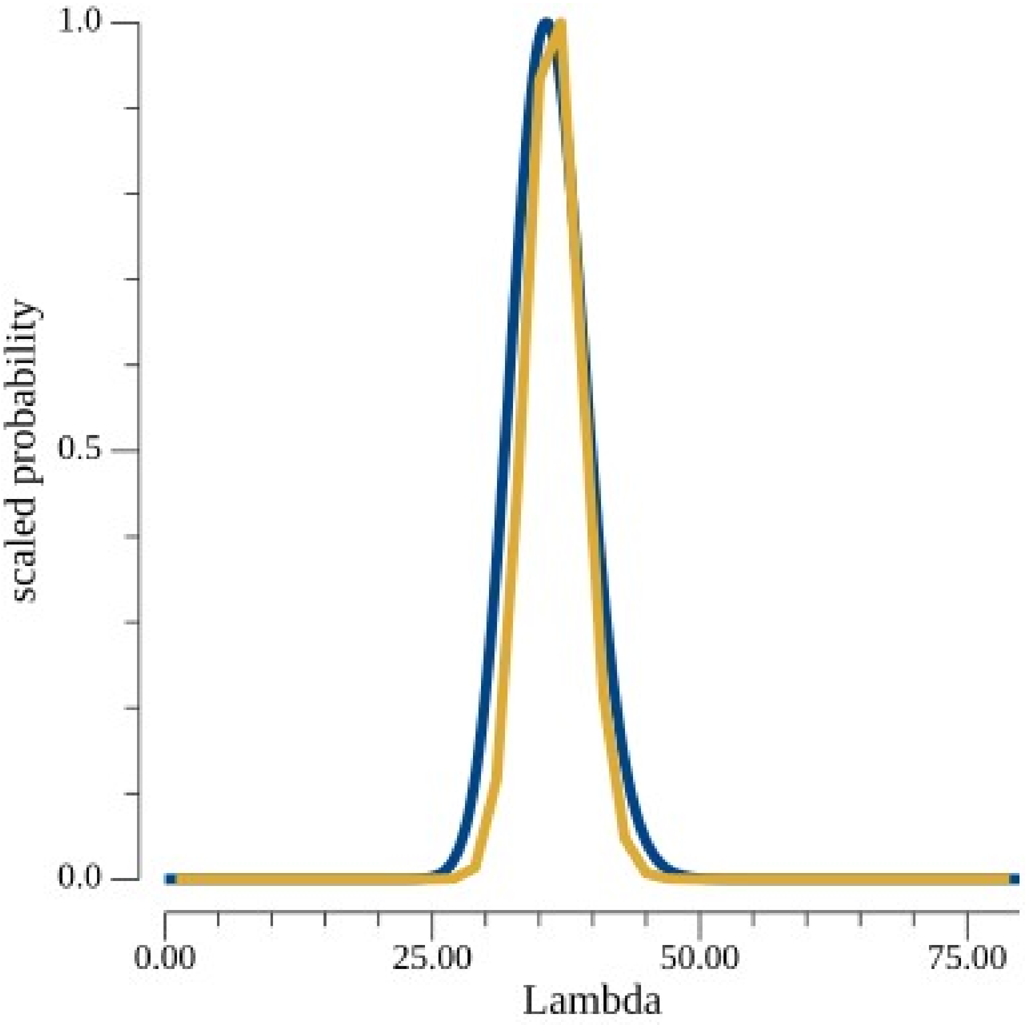


***Supplementary figure 3**. In yellow, the estimated posterior distribution of the λ parameter (using a flat uniform prior) for the Sapindaceae data set; in blue, the gamma distribution (α = 108.0 and β = 3.0) used to approximate the posterior distribution.*

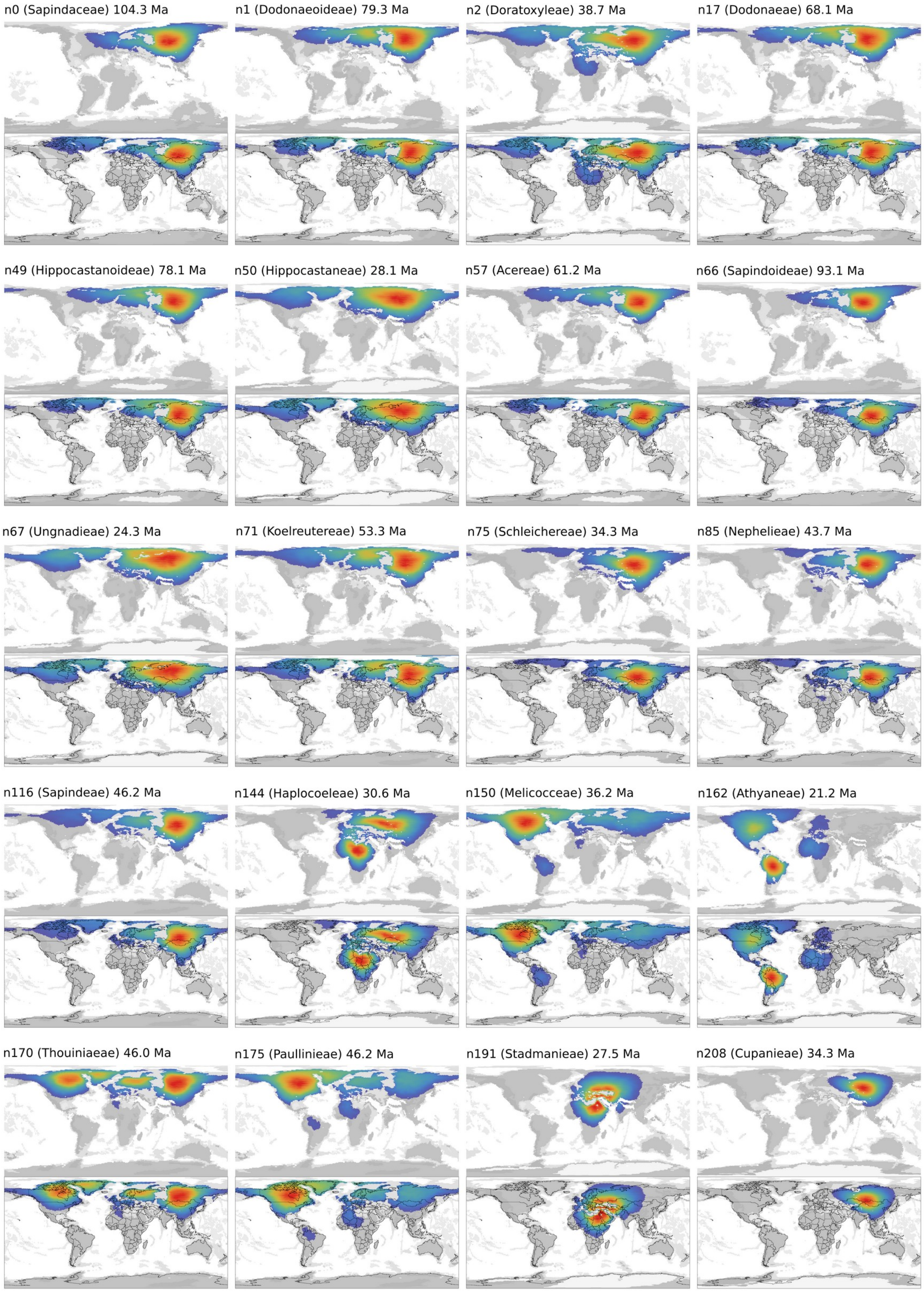
n0 (Sapindaceae) 104.3 Ma
n1 (Dodonaeoideae) 79.3 Ma
n2 (Doratoxyleae) 38.7 Ma
n17 (Dodonaeae) 68.1 Ma
n49 (Hippocastanoideae) 78.1 Ma
n50 (Hippocastaneae) 28.1 Ma
n57 (Acereae) 61.2 Ma
n66 (Sapindoideae) 93.1 Ma
n67 (Ungnadieae) 24.3 Ma
n71 (Koelreutereae) 53.3 Ma
n75 (Schleichereae) 34.3 Ma
n85 (Nephelieae) 43.7 Ma
n116 (Sapindeae) 46.2 Ma
n144 (Haplocoeleae) 30.6 Ma
n150 (Melicocceae) 36.2 Ma
n162 (Athyaneae) 21.2 Ma
n170 (Thouiniaeae) 46.0 Ma
n175 (Paullinieae) 46.2 Ma
n191 (Stadmanieae) 27.5 Ma
n208 (Cupanieae) 34.3 Ma

***Supplementary figure 4**. Pixel posterior probabilities for the nodes assigned to the family, subfamily, and tribal ranks. Monogeneric groups are not displayed. For the node IDs, see supplementary figure 3.*
